# Validation of a Mathematical Model of Cancer Incorporating Spontaneous and Induced Evolution to Drug Resistance

**DOI:** 10.1101/2019.12.27.889444

**Authors:** Jana L. Gevertz, James M. Greene, Eduardo D. Sontag

## Abstract

This paper continues the study of a model which was introduced in earlier work by the authors to study spontaneous and induced evolution to drug resistance under chemotherapy. The model is fit to existing experimental data, and is then validated on additional data that had not been used when fitting. In addition, an optimal control problem is studied numerically.

## 1 Introduction

The ability of cells and tissues to alter their response to chemical and radiological agents is a major impediment to the success of therapy. Such altered response to treatment occurs in infections by bacteria, viruses, fungi, and other pathogens, as well as in cancer [1, 2, 3, 4]. Despite advances in recent decades, this reduction in the effectiveness of treatments, broadly termed *drug resistance*, remains poorly understood, and in some circumstances is thought to be inevitable [5].

One of the most clinically important examples of drug resistance is that involving the treatment of cancer via chemotherapies. Drug resistance is a primary cause of treatment failure, with a variety of molecular and microenvironmental causes already identified [6]. For example, the upregulation of drug efflux transporters, enhanced DNA repair mechanisms, modification of drug targets, stem cells, irregular tumor vasculature, environmental pH, immune cell infiltration and activation, and hypoxia have all been identified as mechanisms which may inhibit treatment efficacy [7, 8, 6, 9, 4, 10, 11, 12, 13, 6, 14]. Experimental and mathematical research continues to shed light on drug resistance, but many challenges remain.

Aside from understanding the mechanisms by which resistance to therapies may manifest, a fundamental question is *when* resistance arises. With respect to the initiation of therapy, resistance can be classified as either *pre-existing* or *acquired* [6]. *The term pre-existing (also known as intrinsic*) drug resistance is reserved for the case when the organism contains a sub-population (or a tumor contains a sub-clone) which resists treatment prior to the application of the external agent. Examples of the presence of extant resistance inhibiting treatments are abundant in bacteria and cancer, including genes originating from phyla Bacteroidetes and Firmicutes in the human gut biome [15], BCR-ABL kinase domain mutations in chronic myeloid leukemia [16, 17], and MEK1 mutations in melanoma [18]. Conversely, acquired resistance describes the phenomenon in which resistance first arises during the course of therapy from an initially drug-sensitive population. The question of whether resistance is pre-existing or acquired is a classical one in the context of bacterial resistance to a phage [19].

The study of acquired resistance is complicated by the question of *how* resistance emerges. Resistance can be *spontaneously* (also called *randomly*) acquired during treatment as a result of random genetic mutations or stochastic non-genetic phenotype switching. These cells can then be selected for in a classic Darwinian fashion [20]. Resistance can also be *induced* (or *caused*) by the presence of the drug [21, 22, 23, 20, 24, 25, 26]. That is, the drug itself may promote, in a “Lamarckian sense”, the (sometimes reversible) formation of resistant cancer cells so that treatment has contradictory effects: it eliminates cells while simultaneously upregulating the resistant phenotype, often from the same initially-sensitive (*wild-type*) cells.

The fundamentally distinct scenarios in which resistance may arise are illustrated in Fig. 1. Note that in each of the three cases (pre-existing, spontaneous acquired, and induced acquired), the post-treatment results are identical: the resistant phenotype dominates the microenvironment. However, the manner by which resistance is generated in each case is fundamentally different, and the transient dynamics (both drug-independent and drug-dependent) may vary drastically in each scenario, so that therapeutic design may have significant effects on clinical outcome.

**Figure 1:**
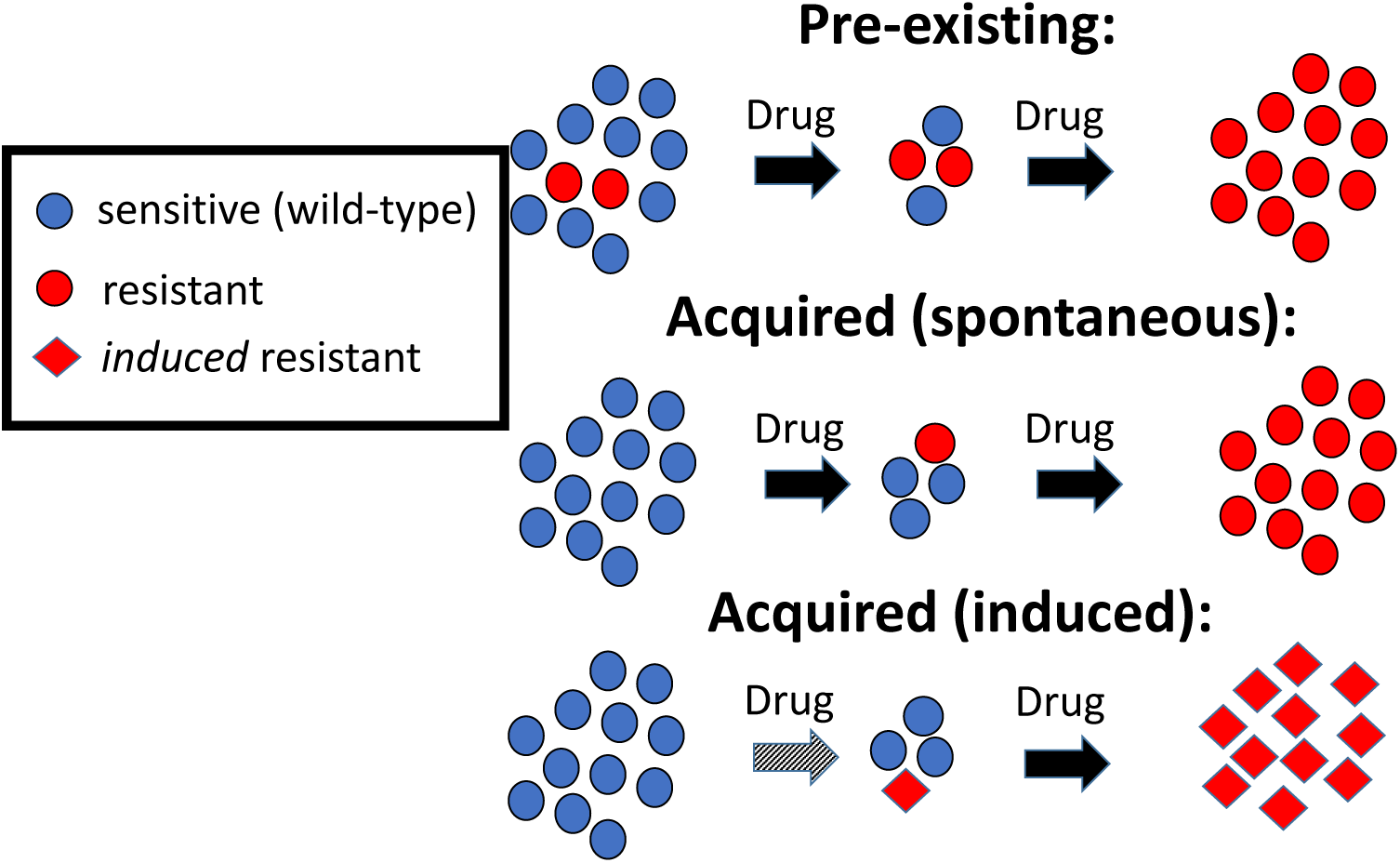
Three distinct ways in which drug resistance may be generated. In the first row, resistance is pre-existing, so that resistant clones are selected during therapy. In spontaneously acquired resistance, random genetic and epigenetic modifications produce a resistant phenotype during therapy, where again selection dominates. In the case of induced acquired resistance, the application of the drug *causes* the resistant mutant to arise, so that the mechanism is *not* simply random variation (e.g. mutations) followed by classical Darwinian selection. Instead, there is a “Lamarckian” component (grey arrow). Note that any combination of these three mechanisms may be causing resistance in a single patient.

As discussed above, there is experimental evidence for these three forms of drug resistance. However, differentiating them experimentally is non-trivial. For example, what appears to be drug-induced acquired resistance may simply be the rapid selection of a very small number of pre-existing resistant cells, or the selection of cells that spontaneously acquired resistance [27, 20]. It is in this realm that mathematics can provide invaluable assistance. Formulating and analyzing precise mathematical models describing the previously mentioned origins of drug resistance can lead to novel conclusions that may be difficult, or even impossible given current sequencing technology, to determine utilizing experimental methods alone. The main goal of this work is to mathematically study *when* and *how* drug resistance develops, and its implications for therapeutic design and treatment outcome. The paper is organized as follows. In Section 2 we will focus on a particular set of experimental data from the Huang lab which seeks to answer the questions of when and how drug resistance develops [20], as well as a prior mathematical model that was motivated by this work [28]. In Section 3 we turn to a novel mathematical model to describe the data in [20], and describe the methodology we utilize to fit the data. We also introduce the optimal control problem that is numerically investigated in the subsequent section. In Section 4 we demonstrate that our newly-proposed model not only gives excellent fits to a training data set, but also very well describes a validation data set. Therefore our numerical optimal control could represent an improved way to administer a cancer drug that can induce resistance. In Section 5, we end with some concluding remarks and directions for future work.

## 2 Prior Experimental and Modeling Work

While there is growing experimental evidence for the *induction* of resistance, herein we will focus on work from the Huang lab [20]. In this work, the relative contribution of selection and induction of drug resistance was assessed in HL60 leukemic cells treated with the chemotherapeutic agent vincristine. Using the expression of the MDR1 gene as a measure of drug resistance, it was found that the expression of MDR1 was mediated by cell-individual induction, and not by the selection of cells with pre-existing expression of MDR1. In other words, the study concluded that the phenotypic plasticity of HL60 cancer cells allows these cells to temporarily change their gene expression to a more resistant state in response to treatment-related stress [20, 24].

The experiments in [20] provide a wealth of data about cancer evolution in response to varying doses of vincristine, though much of the fitting and validation done herein will rely on the dataset shown in Fig. 2. This data describes the effective growth curves (cell number as a function of time) of two cell subpopulations either with or without treatment: 1) an MDR1^High^ subpopulation which is assumed to be drug-resistant, and 2) an MDR1^Low^ subpopulation which is assumed to be drug-sensitive.

**Figure 2:**
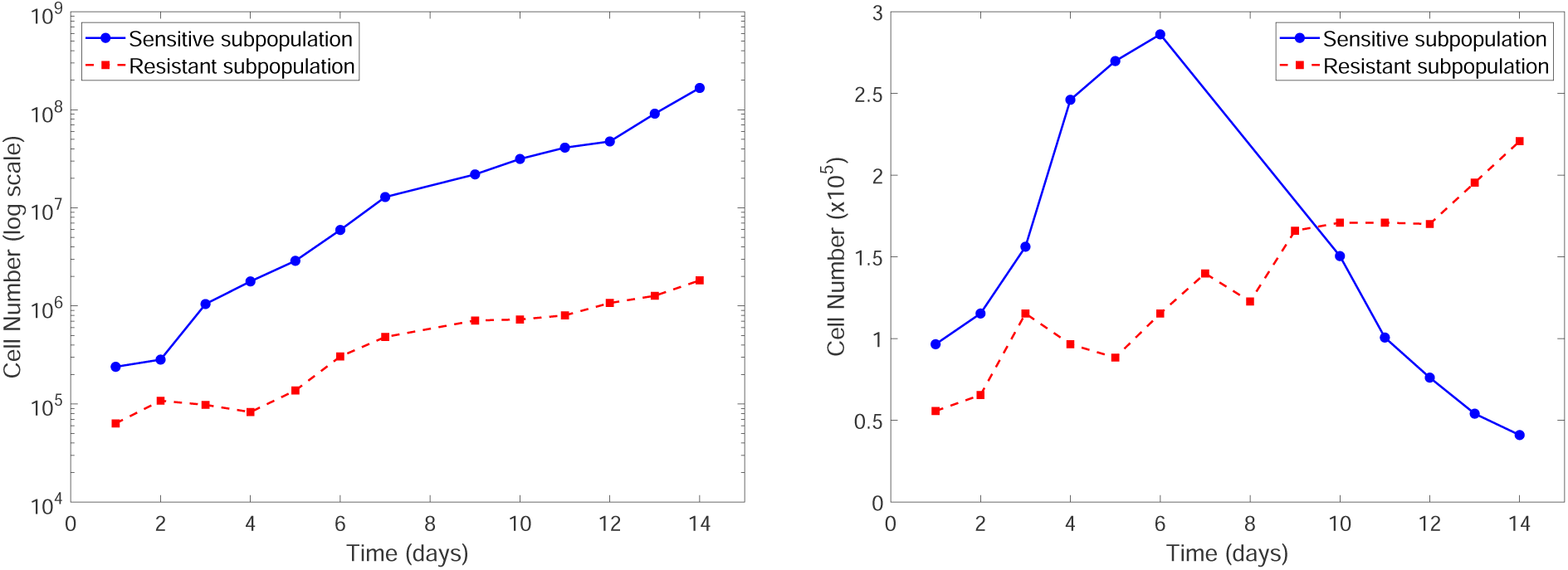
Time course data [20] showing the change in cell number for an initially sensitive subpopulation of HL60 cells (solid blue curve), and an initially resistant subpopulation (dashed red curve). Dynamics in the absence of a drug challenge are shown on the left, and in the presence of vincristine are shown on the right. Due to probable experimental error (personal communication from the authors of [20]), we are withholding the data at days 8 and 9 when the sensitive subpopulation is treated with vincristine.

There have been many efforts to further our understanding of drug-resistance in cancer using mathematical models (see [29, 30] for reviews), though only a small percent of these have focused on resistance induction. Of those models that consider drug-induced resistance, many are dose-independent models that implicitly model the effects of the drug [20, 31, 32, 33, 34]. Models which explicitly incorporate the drug, as we do here, have also been developed in [35, 36, 37]. To our knowledge, none of these previously-developed models simultaneously studied the impact of selection of pre-existing resistant cells along with the induction of acquired resistance (though work in [35, 38, 39] did consider each of these phenomena independently). In an effort to better understand the role of selection and induction in the evolution to drug resistance, in [28] we introduced a simple, phenomenological mathematical framework to distinguish between spontaneous and induced resistance. Specifically, we considered a tumor population consisting of sensitive (*S*), non-reversible resistant (*R*_*n*_), and reversible resistant (*R*_*r*_) cells, whose dynamics are described by the following set of ordinary differential equations (ODEs):

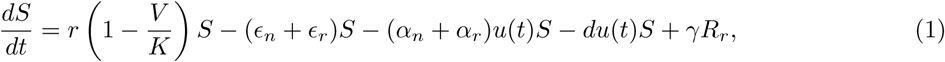

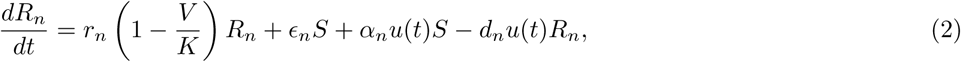

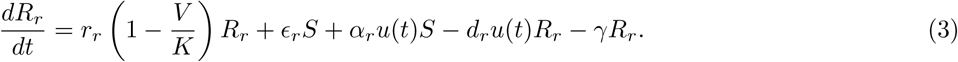

Here *u*(*t*) represents the effective applied drug dosage at time *t*, and is thought of as a control, and *V* is the total tumor population (*V* = *S* + *R*_*n*_ + *R*_*r*_). By definition, the drug is less effective against resistant cells, so we assume that *d*_*n*_, *d*_*r*_ < *d*. We also assume that *r* > *r*_*n*_, *r*_*r*_, as experimental evidence supports that resistant cells grow slower than nonresistant cells [40, 41]. All cells compete for resources via a joint carrying capacity *K*, and resistance may develop in *both* a spontaneous drug-independent (*ϵ*_*n*_*S, ϵ*_*r*_*S*) and induced drug-dependent (*α*_*n*_*u*(*t*)*S, α*_*r*_*u*(*t*)*S*) manner. In eqns. (1) - (3), non-reversible cells *R*_*n*_ can be thought of as resistant cells that arise via genetic mutations - as “undoing” genomic mutations is unlikely, reversibility is assumed negligible. However, resistant cells *R*_*r*_ may form via phenotype-switching, which is often reversible [23, 20], so that we included a re-sensitization term (*γR*_*r*_) as well. The drug induction terms *α*_*n*_*u*(*t*)*S* and *α*_*r*_*u*(*t*)*S* assume a log-kill hypothesis in which the rate of induced resistance is proportional to the applied dosage.

We have previously studied the effects of resistance induction in a simplified version of this model, where we assumed that the reversible and non-reversible resistant cells exhibit identical kinetics (*R* := *R*_*n*_ = *R*_*r*_, *ϵ* := *ϵ*_*n*_ = *ϵ*_*r*_, *α* := *α*_*n*_ = *α*_*r*_, *r*_*n*_ = *r*_*r*_, *d*_*n*_ = *d*_*r*_):

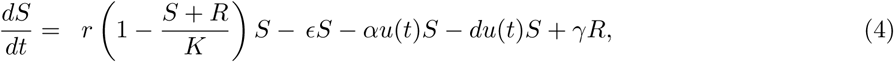

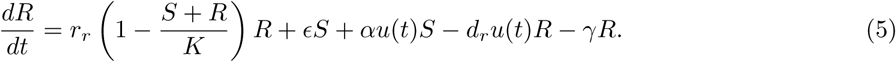

Using this simplified model, we found a *qualitative* distinction in tumor response to two standard treatment schedules (continuous versus pulsed) when the drug has the same cytotoxic potential but different levels of induction (i.e. same *d* but different *α* in eqns. (4) - (5)). In particular, for a fixed set of patient-specific parameters, when a drug cannot induce resistance, constant therapy significantly outperforms pulsed therapy, as the time to treatment failure is nearly seven times longer using constant therapy. However, when a drug can induce resistance, pulsed therapy improves the duration of therapeutic efficacy when compared to constant treatment [28]. This illustrates how the mechanism of drug resistance can strongly influence response to a treatment protocol. A rigorous analysis of the optimal control structure (both theoretical and numerical) can be also be found in [42].

## 3 Mathematical Model and Methods

In Section 4, we will argue that the originally proposed system in eqns. (4)-(5) cannot adequately describe the experimental data in [20]. Herein, we propose an alternative mathematical model for distinguishing between spontaneous versus induced evolution to drug resistance. We also discuss the methodology used to fit the model to the experimental data in the presence of vincristine (right plot in Fig. 2), and our process for validating the model using the experimental data in the absence of treatment (left plot in Fig. 2). We conclude this section with a description of the numerical optimal control methodology.

### 3.1 Revised Mathematical Model

The initial increase in the number of cells when the sensitive cell subpopulation is treated with vincristine (see blue curve in right-hand plot in Fig. 2) suggests that the drug does not immediately cause a decrease in the sensitive cell population. This observation led us to to develop a modified version of our model that directly incorporates drug concentration *y*, instead of having an indirect measure of the drug through the control *u*. We assume that there is a “delay” in drug action, likely due to cell cycle phase dependence. This is a reasonable assumption for vincristine, as it partially works by halting the formation of microtubules needed for cells to separate during metaphase. This delay eventually results in the apoptosis of the affected cell [43].

Although a carrying capacity was built into our model in eqns. (4)-(5), here we use simple exponential growth. This is because our original model was envisioning an *in vivo* setting, whereas herein we are modeling *in vitro* data under non-limiting growth conditions, as shown by the growth data in Fig. 2 which exhibits a clear exponential, non-saturated growth. We propose the following modified model:

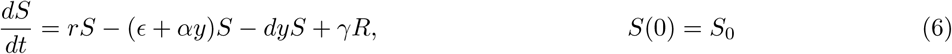

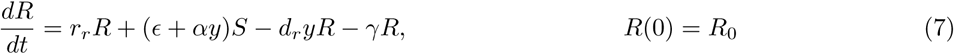

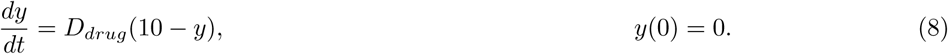

Note that, if we changed the exponential terms to logistic terms, this model is equivalent to the original model in eqns. (4)-(5) in the absence of drug (that is, when *α* = *d* = *d*_*r*_ = *D*_*drug*_ = 0). The coefficient 10 is used in eqn. (8) to represent the applied dosage (in *nM*) of vincristine in Fig. 2. See Section 3.3 for a control formulation with arbitrary dose *u*.

### 3.2 Fitting Methodology

Our goal here is to fit the model in eqns. (6)-(8) to the treatment data in Fig. 2. The no-drug data will be withheld from the fitting, to be subsequently used for validation purposes. The model has eight parameters that should be shared across the two treatment datasets, meaning they should be unaffected by whether we start with a sensitive or resistant subpopulation. In the experiment that starts with a sensitive subpopulation, *R*(0) must be zero, but *S*(0) can be treated as a free parameter. Similarly, in the experiment that starts with a resistant subpopulation, *S*(0) must be zero, but *R*(0) can be treated as a free parameter. As previously discussed, we assume that all parameters are non-negative, that sensitive cells grow faster than resistant cells (*r > r*_*r*_), and that the drug is more effective at killing sensitive cells (*d > d*_*r*_).

To fit our model to the experimental data, we begin with a quasi Monte Carlo sampling technique that uses Sobol’s Low Discrepancy Sequences (LDS) [44] to identify a neighborhood in 10D-parameter space where a global optimum should occur. Here, 10D-parameter space is defined by {*r, r*_*r*_, *ϵ, α, d, d*_*r*_, *γ, D*_*drug*_, *S*_0_, *R*_0_}. Note that the initially sensitive and initially resistant subpopulations are fit together, meaning we are fitting the parameter set {*r, r*_*r*_, *ϵ, α, d, d*_*r*_, *γ, D*_*drug*_} across both datasets simultaneously. On the other hand, *S*_0_ > 0 is only fit to the initially sensitive population which has *R*(0) = 0, and *R*_0_ > 0 is only fit to the initially resistant population with has *S*(0) = 0. The best-fit parameter set is defined as the set that minimizes the sum of the square error between the model predictions and the data.

After a parameter set has been selected among the randomly sampled sets chosen by Sobol’s LDS, an optimization is carried out in order to minimize the error, starting from this point and refined using simulated annealing. Having observed that the landscape of the objective function near the optimal parameter set does not appear to contain local minima, we randomly perturb the LDS-chosen parameter set, and accept any parameter changes that decrease the value of the objective function. This random perturbation process is repeated until no significant change in the sum of the square error can be achieved, and we call this final parameter set the optimal parameter set.

After finding the optimal parameter set, we again used Sobol’s LDS to sample points in a neighborhood about the optimal parameter set. We tested each sampled parameter set to see if it gives a goodness-of-fit within 10% of the optimal fit. Any such parameter sets are classified as suboptimal sets. These suboptimal parameter sets will be employed as we validate the model, and use the model to make treatment-related predictions.

### 3.3 Numerical Optimal Control

As the authors of [20] consider various constant dosing strategies over a fourteen day period, we formulate an optimal control problem for the parameters describing the data set in Fig. 2. Precisely, we generalize the mathematical model above as follows:

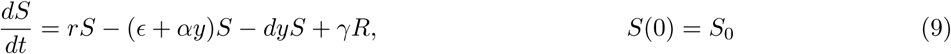

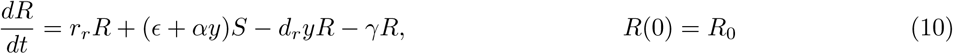

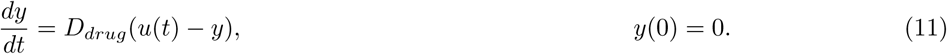

Note that the only difference with the above and eqns. (6)-(11) is that we allow an arbitrary applied background dose *u*(*t*), which we assume as a bounded Lebesgue measurable function. Since the authors in [20] apply vincristine between 0 and 60 *nM*, we assume likewise that

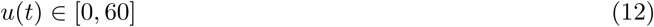

for all *t* ∈ [1, 14]. We thus consider a fixed-time problem, where we minimize the final tumor volume on the fourteenth day of treatment. To account for toxicity of vincristine [45, 46], we also include a penalty term in the below objective corresponding to the total administered dose. Thus, we minimize the following objective:

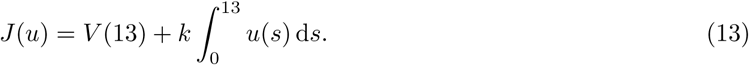

Here *k* is a constant that is used to trade-off final tumor size and the amount of drug applied; numerical results for various values are provided in Section 4.4. Note that we consider treatment beginning on day 0, as opposed to day 1 as appears in Fig. 2. We thus numerically investigate the control structure *u* = *u*(*t, k*) which minimizes eqn. (13) for the optimal parameter set obtained via the fitting methods discussed in Section 3.2.

## 4 Results & Discussion

### 4.1 Insufficiency of Original Model

Our aim here is to demonstrate that the model in eqns. (4)-(5) (adapted to have exponential growth terms instead of logistic, as explained above) cannot adequately describe the data in Fig. 2. In particular, here we consider:

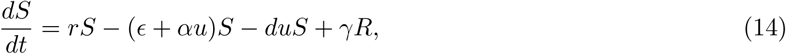

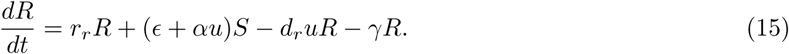

The control *u*(*t*) is now assumed to be constant (*u*(*t*) := *u*), consistent with the experimental setup in [20].

In order for this model to be a sufficient representation of the experimental data, we require that in the presence of drug (*u* ≠ 0), both the initially sensitive subpopulation and the initially resistant subpopulation must initially increase in size (see right-hand portion of Fig. 2). Further, we require that the initially sensitive population eventually decreases during the treatment window. In Result 1, we argue that our model in eqns. (14)-(15) cannot exhibit this behavior subject to a constant dosing regimen.

#### Result 1

*Under the assumption of constant dosage, the system in eqns. (14)-(15) cannot describe the time course data for the evolution of an initially sensitive and resistant subpopulation treated with vincristine*.

*Proof*. Consider the system in eqns. (14)-(15) and assume 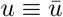 is the constant dosage. Defining

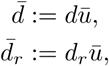

the dynamics for the tumor volume under constant treatment can be described by

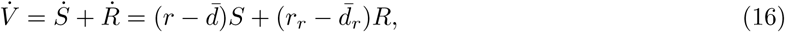

with all parameters positive. In order for the initially sensitive subpopulation to increase in size in the presence of drug, we require that when *S*(0) > 0 and 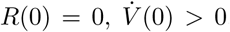. According to eqn. (16), this requires

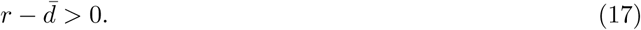

On the other hand, in order for the initially resistant subpopulation to increase in size in the presence of drug, we need that when *S*(0) = 0 and 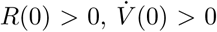. Since 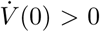 in this case as well, eqn. (16) requires that

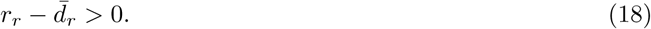

Since these two cases are in the same phase plane (it is the same dose, just different initial conditions), the fact that 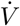 must change sign in the *S*(0) > 0 case means that in the first quadrant, there must be a point at which 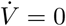. Since eqn. (18) requires that 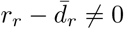, we get

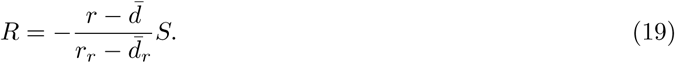

Since we need *R >* 0 when *S >* 0 for this to occur in the first quadrant, we require that 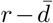 and 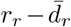 have opposite signs. However in eqns. (17) and (18), we previously argued that 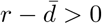 and 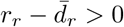. Thus we have reached a contradiction, which proves that our original model in eqns. (14)-(15) cannot accurately describe the time course data in Fig. 2, subject to a constant dosing regimen of vincristine. □

### 4.2 Fitting to Modified Model

Here we employ the previously-described fitting methodology to simultaneously fit the vincristine data in Fig. 2 in the cases of an initially sensitive and an initially resistant subpopulation. The results of our fitting can be seen in Fig. 3. We observe a very good agreement between the model and both datasets. The optimal parameter values, along with the range of values each parameter can take on in the suboptimal parameter set, are shown in Table 1.

**Table 1:** Best-fit parameter values, range for suboptimal parameters, and the absolute value of the maximum deviation for which the parameter set remains within suboptimal set

| Parameter | Best-Fit Value | Suboptimal Range | \|Max Deviation\| |
| --- | --- | --- | --- |
| $r$ | 0.5183 | [0.47681, 0.5858] | 13.0% |
| $\epsilon$ | 0.006913 | [0.001003, 0.01026] | 85.5% |
| $\alpha$ | 0.0002656 | [0.000028, 0.003989] | 1399% |
| $d$ | 0.7244 | [0.3217, 1.386] | 91.2% |
| $\gamma$ | 0.002935 | [0.0001780, 0.03677] | 1153% |
| $r_r$ | 0.1466 | [0.09610, 0.1596] | 34.4% |
| $d_r$ | 0.05823 | [0.002169, 0.09594] | 96.4% |
| $D_{drug}$ | 0.01457 | [0.007425, 0.03194] | 119.1% |
| $S_0$ (initially sensitive) | 77867 | [67221, 87501] | 13.7% |
| $R_0$ (initially resistant) | 62230 | [56027, 73002] | 17.3% |

**Figure 3:**
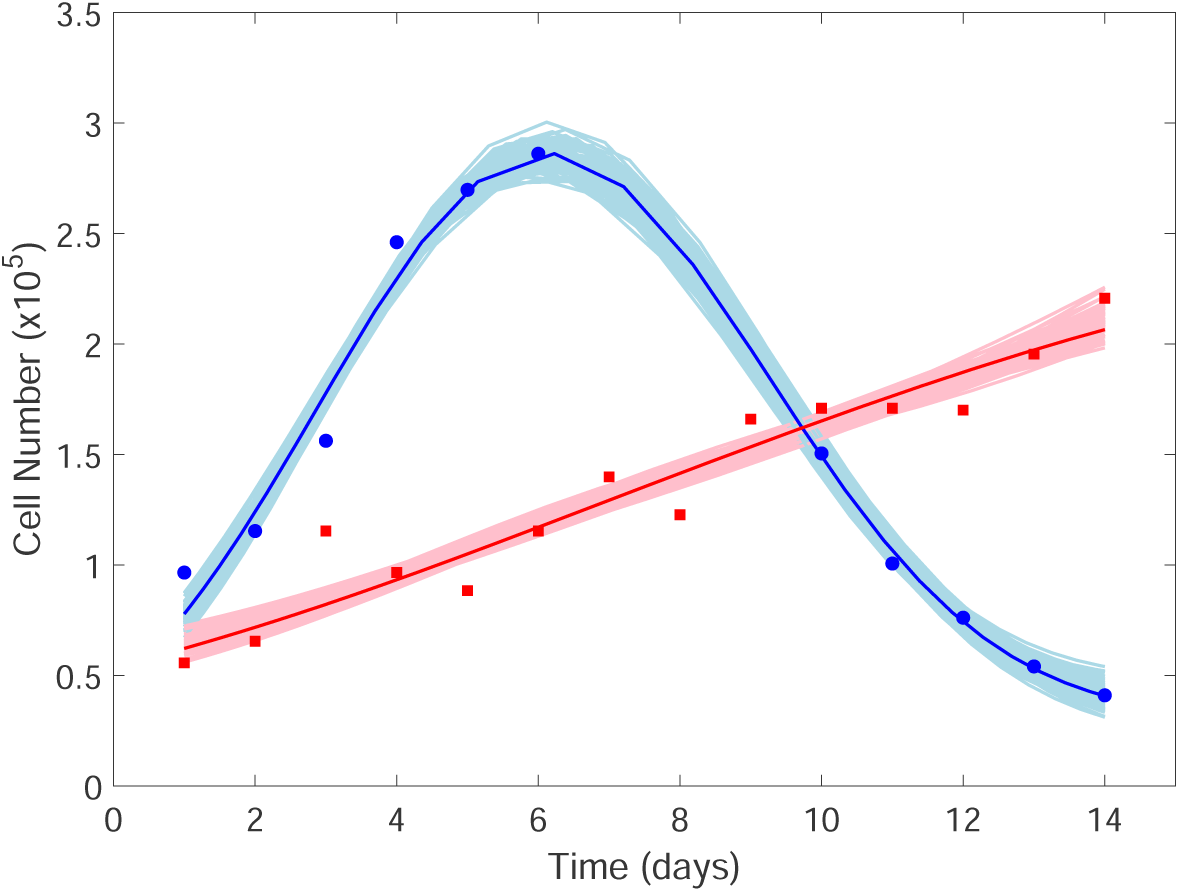
Fits to the data describing tumor size subject to treatment with vincristine [20]. The blue circle data points represent the experimental data when an initially sensitive subpopulation is treated with vincristine. The solid blue curve is the best-fit of the system in eqns. (6)-(8) to that data, and the light blue curves are suboptimal parameter sets. The red square data points represent the experimental data when an initially resistant subpopulation is treated with vincristine. The solid red curve is the best-fit of model to that data, and the light red curves correspond to representative suboptimal parameter sets.

### 4.3 Validation of Modified Model

In an effort to *validate* the predictive abilities of the model, we now consider the no-drug data shown in Fig. 2. We fix all drug-independent parameters as shown in Table 1, and set all drug-related parameters (*α, d, d*_*r*_, *D*_*drug*_) equal to zero. For the initially sensitive subpopulation, the only degree of freedom is the nonzero initial condition *S*_0_. Similarly, for the initially resistant subpopulation, the only degree of freedom is the nonzero initial condition *R*_0_. Each of the free initial conditions is fit to its corresponding dataset using the Nelder-Mead simplex method as implemented by MATLAB’s fminsearch function. As shown in Fig. 4, our model does an excellent job of describing the no-drug data for both the initially sensitive and initially resistant subpopulations.

**Figure 4:**
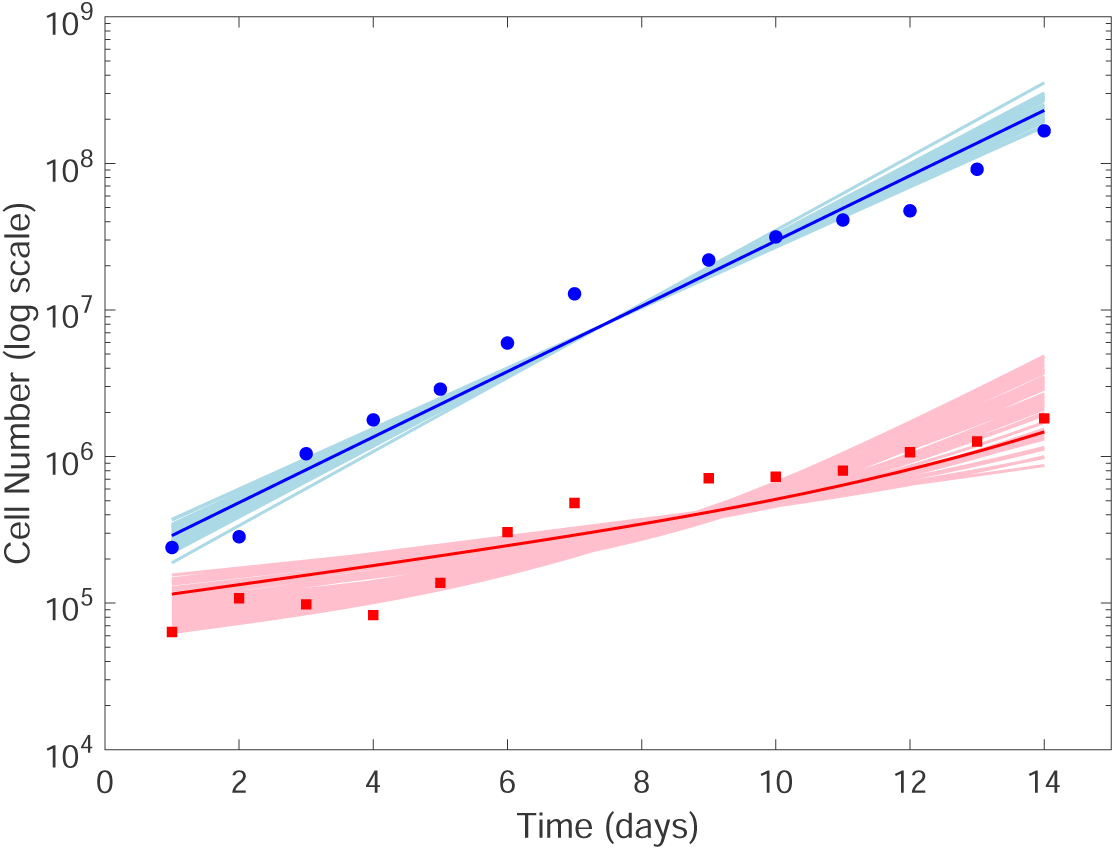
Validation of model against the tumor growth data in the absence of treatment [20]. The blue circle data points represent the experimental data when an initially sensitive subpopulation grows without treatment. The solid blue curve is the model prediction when fitting only the initial amount of sensitive cells (with optimal *S*_0_ = 28901), and the light blue curves are the predicted curves for the suboptimal parameter sets. The red square data points represent the the experimental data when an initially resistant subpopulation grows without treatment. The solid red curve is the model prediction when fitting only the initial amount of resistant cells (with optimal *R*_0_ = 11535), and the light red curves are the predicted curves for the suboptimal parameter sets.

As another way to validate the model, we turn to cell-sorting experiments in [20] in which HL60 cells are sorted based on MDR1 expression levels. It was found that, at steady-state, 97.8% of the cell population can be classified as sensitive. To determine the steady-state fraction of sensitive cells predicted by our model, we derived the differential equation for the *fraction* of sensitive cells from eqns. (6)-(7) in the absence of drug.

Defining the sensitive fraction as

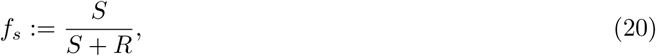

we can derive the following ordinary differential equation for *f*_*s*_:

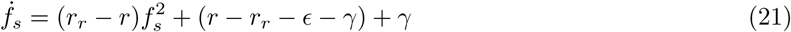

Note that *R/S* + *R* = 1 − *f*_*s*_. The above then has the unique nonnegative globally asymptotically stable steady-state fraction of sensitive cells 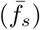 is given by:

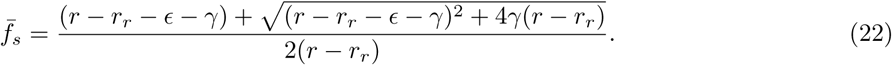

At the optimal parameter set shown in Table 1, we find that 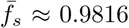, which is in excellent agreement with the experimentally-observed steady-state fraction of 0.978.

For further model validation, we next consider the repopulation experiments in [20]. In these experiments, a population of HL60 cells at its steady-state distribution of MDR1 expression were sorted into a sensitive and resistant subpopulation. Each subpopulation was allowed to grow in the absence of drug until the steady-state distribution of MDR1 in the original population was recovered. To use this dataset to validate our model, we need to know the initial number of cells used in the repopulation experiment. However, the experimental protocol only considered the initial *fraction* of sensitive and resistant cells in the repopulated pools. However, note that the steady-state fraction (22) is insensitive to the raw initial numbers provided we start with a nonzero fraction of both sensitive and resistant cells.

We exploit this knowledge for the sensitive cell repopulation experiment by noting that our numerical simulations indicated that if we start with a sensitive fraction greater than 0.9996 but less than 1 (so there is no dependence on the raw initial number of sensitive cells), the result is that 99.7% of the population is sensitive after 12 hours (see Table 2). This is compared to the 97.8% observed experimentally [20]. For the resistant cell repopulation experiment, the experimental data in [20] indicates that we start with a resistant subpopulation, four days later the population is composed of 48.3% sensitive cells. Using this data as the initial condition for modeling the resistant cell repopulation experiment, we find that that the model predicts that, after 17 days, 97.3% of the cells are sensitive (see Table 2) compared with the 97.8% that is observed in the experimental data [20]. There is fairly good agreement with the experimental data, if one takes the cut-off value for resistance as shown in [20] that corresponds to 97.8% resistant cells. This shows a remarkable *predictive* ability of our model, which lumped a complicated distribution into just two classes and was fit to aggregate, not single-cell, data.

**Table 2:**
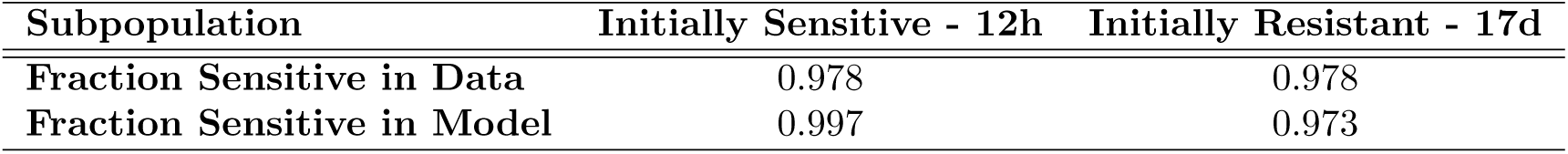
Comparing experimental data and model predictions for the repopulation experiments. The initially sensitive subpopulation is compared at 12 hours, the time it took to recover the baseline distribution in the experimental data. The initially resistant subpopulation is compared at 17 days, the time it took to recover the baseline distribution in the experimental data.

### 4.4 Numerical Optimal Control

Using parameters found in Section 4.2 (Table 1), we investigate the optimal control problem introduced in Section 3.3. For the following numerical computations, we re-scale eqns. (9)-(11) by the initial sensitive population *S*_0_, so that we consider the following initial conditions:

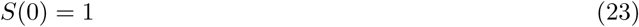

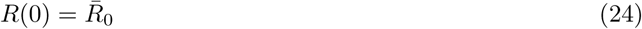

For the remainder of the section, we assume that treatment begins at the equilibrium distribution discussed in [20], so that

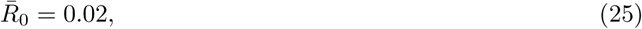

and numerically solve the optimal control problem numerically using the GPOPS-II MATLAB sparse non-linear programming solver [47].

We begin by letting *k* = 0 in eqn. (13), so that the toxicity effects of vincristine are disregarded in the dosing strategy. Numerical results appear in Fig. 5. Note that the applied optimal control *u*(*t*) is constant at the maximum allowed dose of 60 *nM*, and this protocol appears to cause the eradication of the entire tumor (including the resistant subpopulation).

**Figure 5:**
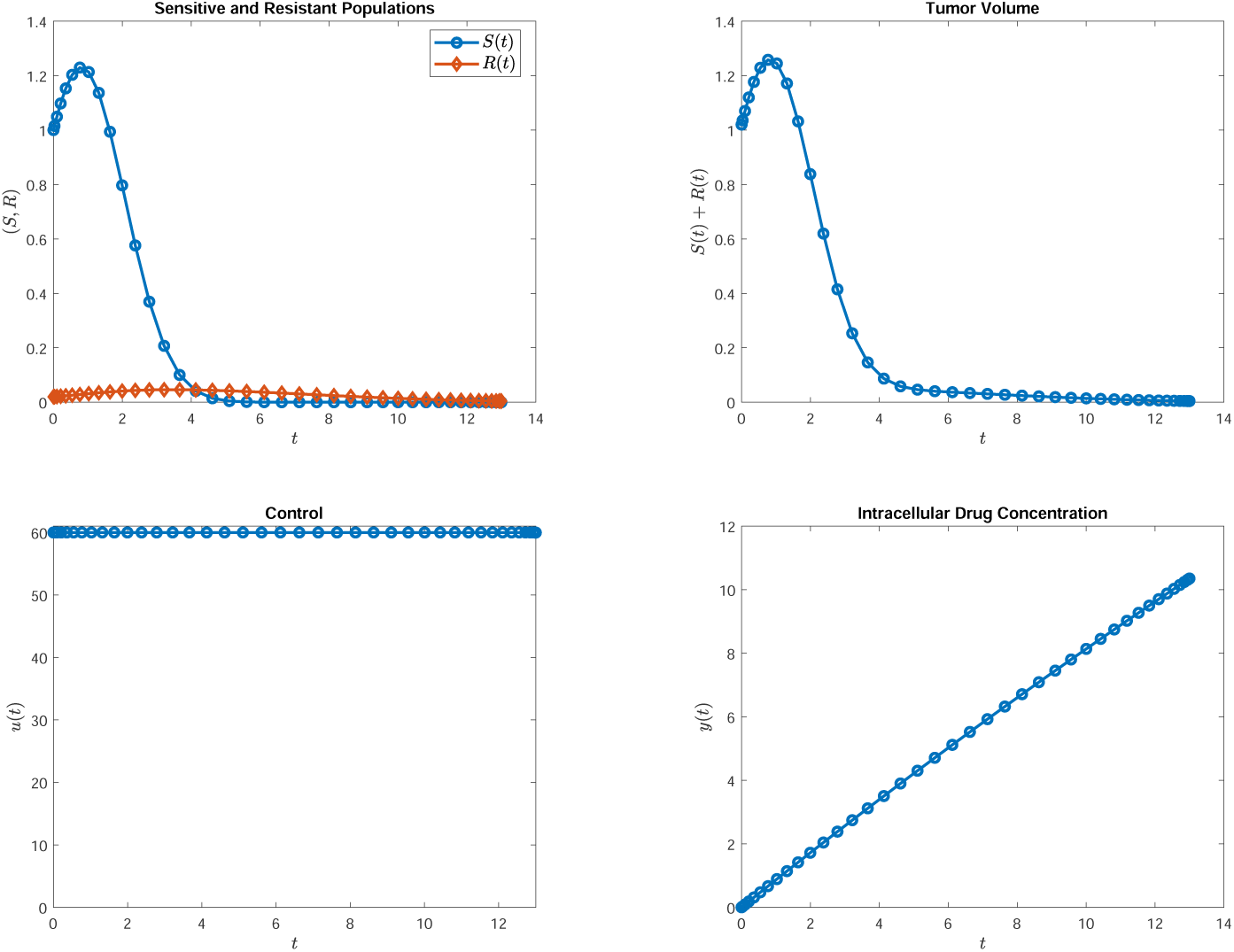
Optimal control for the case when *k* = 0 in eqn. (13). Top left: sensitive and resistant compartments. Top right: tumor volume. Bottom left: control *u*. Bottom right: absorbed drug concentration *y*. Note that the computed optimal control is constant at the maximally applied dosage of 60 *nM*.

We next investigate how the control strategy changes as toxicity is considered. We begin by setting the relative weighting constant at *k* = 0.1; results are presented in Fig. 6. Here we see that the treatment begins at the maximal dose of 60 *nM*, but after approximately one day, treatment is turned off due to toxicity effects. Of course, this therapy is less effective in terms of the overall tumor volume at the end of therapy, compared with the treatment presented in Fig. 5. Continuing in this manner (increasing *k*), we observe a similar phenomenon: an initial maximal dosage applied, and it is switched off at some time and remains at zero thereon. Furthermore, this critical switching time appears to increase as *k* increases. This result can be explained heuristically from our model, as the intracellular drug concentration decreases relatively slowly once therapy is removed (see the *y*(*t*) plots, and note that *D*_*drug*_ = 0.01457), so initially applying a large drug dose will allow the effective dose *y* high to remain high, even after *u* is removed. Although not very interesting, other scenarios are being ruled out by the optimization, such as a sequence of pulses whose total area under the curve remain the same.

**Figure 6:**
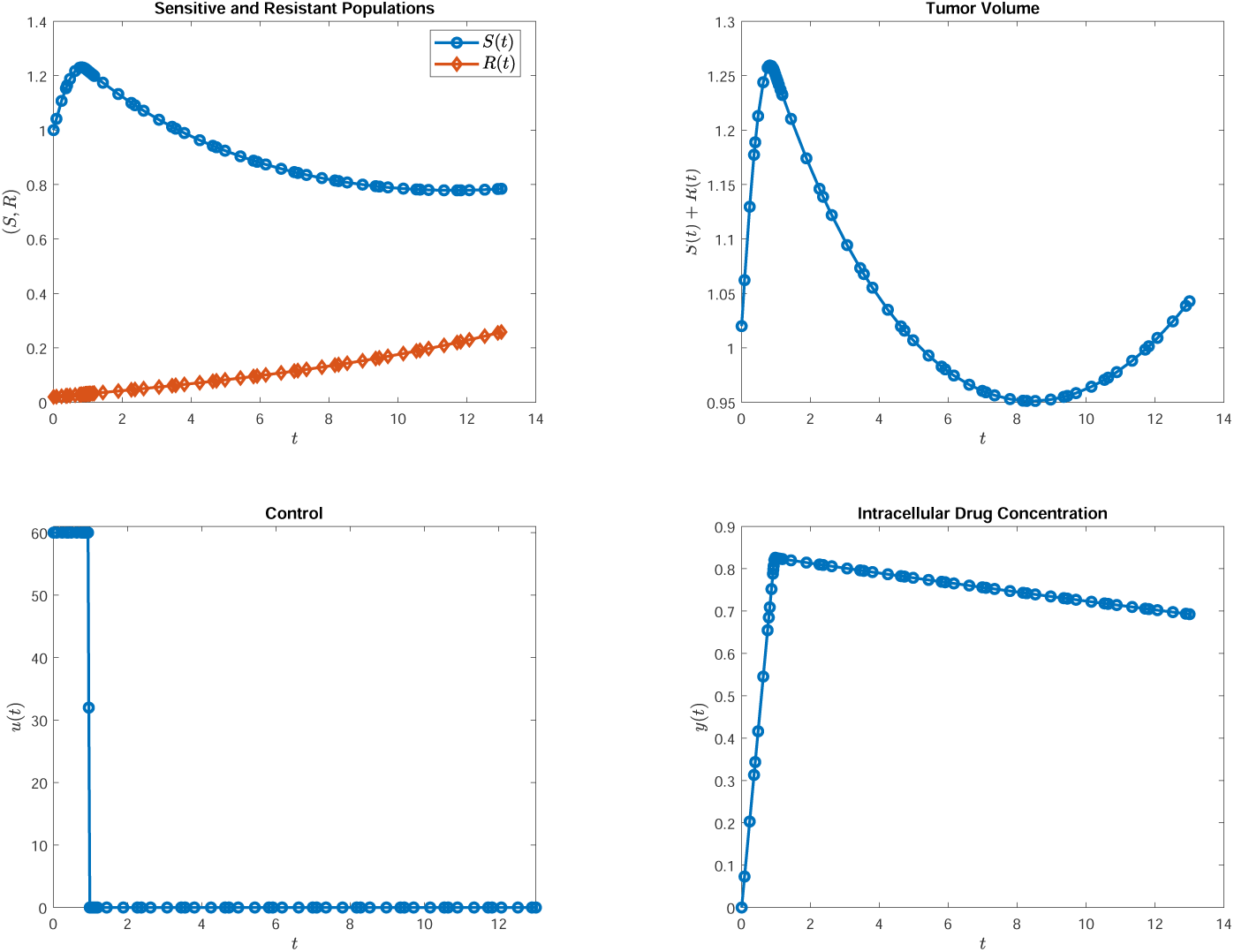
Optimal control and corresponding dynamics for the case when *k* = 0.1 in eqn. (13). See Fig. 5 for corresponding subplots. Note that the control is bang-bang, and switches from *u*_max_ = 60 to *u* = 0 approximately after the first day of treatment.

## 5 Conclusions and Future Work

In this work we have extended a previously introduced model of spontaneous and induced drug resistance which better describes the data presented in [20]. Specifically, we introduced an additional term to model delays in the chemotherapy action due to the cell cycle phase dependence of vincristine. We simultaneously fit this updated model to two experiments conducted in the presence of treatment: an initial population composed entirely of sensitive (MDR1^Low^) cells, and an initial population composed of entirely resistant (MDR1^High^) cells. Although fitting only a small number of parameters (ten), we found very good agreement between the model and both sets of data (Fig. 3). To further validate the model, we compared predictions from our mathematical model to experiments done in the absence of any applied chemotherapy. We again see very close agreement, both in comparison with tumor growth data (Fig. 4), and in re-population experiments (Table 2). Lastly, we used the computed parameter sets to investigate the structure of the optimal drug-dosage regime on a fourteen day treatment window. Here our goal was to balance the final cell population size with the toxicity of vincristine (see eqn. (13)). Our preliminary numerical results suggest a bang-bang structure for the control, where the chemotherapy is initially applied at its maximal dosage, and subsequently turned off for the remainder of the cycle.

Although we have obtained close agreement with our mathematical model and experimental data, the work from the Huang lab [20] contains a large amount of dose-response flow cytometry (FACS) data, which we have obtained directly. Incorporating this data into our modeling framework would allow for further validation, as well as understanding more thoroughly the role induction plays in their experimental system. We are also undertaking a more rigorous analysis of the optimal control structure, as the results presented here are preliminary. Specifically, we are using the Maximum Principle and Lie-algebraic techniques to understand the theoretical structure of the optimal control as parameters vary over the computed suboptimal set. We will also investigate the numerical structure as the treatment window is varied, and investigate how the trade-off between final tumor size and toxicity affects the computed control.

